# Monkeys integrate facial expressions and direct gaze to modulate gaze-following behavior

**DOI:** 10.1101/2025.05.18.654731

**Authors:** Ian Chong, Peter Thier

## Abstract

For humans, being looked at directly can boost our readiness to follow another’s gaze, but can monkeys invite an observer to engage with them in the same way? We trained three rhesus macaques on a head gaze-following task in which the portrait of a demonstrator monkey would face the viewer before turning to look at a distinct spatial target. The demonstrator could look at the viewer with his eyes opened or closed and display different facial expressions. Unlike in humans, we found that direct gaze alone, devoid of an accompanying specific expression failed to influence the latency of the subsequent gaze-following response. However, when combined with threat, direct gaze significantly accelerated gaze-following. In a second experiment, we show that once turned away the expression associated with prior direct gaze no longer mattered; instead a submissive facial expression accompanying the gaze shift delayed gaze-following. Direct comparison of both experiments reveals that expressions accompanying direct gaze trigger earlier gaze-following responses than the same expressions joining gaze aversion. These results document the pronounced behavioral importance of the valuation of expressions signaling danger, arguably creating heightened alertness in monkey observers, thereby priming their gaze-following to allow for immediate conflict resolution.

## 1. Introduction

Humans are able to rely on many forms of nonverbal communication, which include facial expressions, gestures, body poses or gaze to communicate their desires and intentions. When the eyes of the other are gazing directly at an observer – a dyadic interaction usually referred to as “direct gaze” – they signal the other’s intention to approach or interact with the observer (1,2). However, direct gaze may also serve as a means to intimidate and exert dominance, thereby maintaining distance and discouraging approach. Hence, direct gaze can elicit strong arousal, and depending on the social context, may evoke pleasurable or discomforting experiences (3,4). An important potential consequence of direct gaze, drawing the observer’s attention to the eyes of the other is to follow his/her gaze, in case the other may decide to attend to a novel object of interest. By following the other’s gaze the observer becomes able to identify this object and to establish a joint attentional focus on it (5–7). In humans, it has been found that prior direct gaze from a human avatar elicits earlier subsequent gaze-following responses (8). This effect is apparent even though the face avatar had neither identity nor expressions and came without semantic context. Hence, in humans, direct gaze alone is powerful enough to prime us to follow the gaze of the other more readily.

Rhesus macaques are also capable of gaze-following, which is recognized to be a fast, accurate, and highly reflexive process that can be subject to cognitive control, sharing many similarities with human gaze-following (9,10). However, the manner in which they put this into practice is still poorly understood. It is unknown if macaques can use direct gaze to initiate gaze-following like in humans. However, as a relatively despotic species rhesus macaques may shy away from prolonged gaze as it can signal threat and aggression (11). Meanwhile, regardless of clear species-dependent differences in the semantics of facial expressions, nonhuman primates too rely on the meaning of facial expressions in order to guide their interactions with conspecifics. For instance, facial expressions can facilitate gaze-following in Barbary macaques, which exhibit a ‘commenting expression’. This is produced when observing others and has been found to promote gaze-following, informing an observing monkey of further interactions taking place in the vicinity (12). This partially answers how a bout of gaze-following may be kick-started but is still reliant on an observer witnessing the commenting gesture by chance. Moreover, rhesus macaques do not possess a commenting gesture. Other work in long-tailed macaques reported that fear expressions accompanying a gaze-shift can boost gaze-following (13). Yet, the problem of this work and others is that it studied monkey observers reacting to human demonstrators (14,15), and monkeys have been found to not treat human and conspecific facial information in the same way (16).

Champ and colleagues (17) explored behavioral reactions by having a rhesus macaque observer watch two videos displayed side-by-side, showing a dominant and a submissive monkey respectively, in an attempt to simulate two monkeys interacting. They reported an increase in the frequency of gaze-following responses in the interval following the meeting of the eyes. Furthermore, it seemed that submissive expressions of the subordinate monkey increased the number of exploratory saccades made by the observing monkey that landed on the dominant monkey. This suggests that submissive facial expressions can potentially encourage gaze-following behavior due to the need to seek out the source of threat. However, because the observer passively watched these interactions, it is unlikely that they felt part of the interaction, and therefore it is hard to predict the consequences of any coincidental direct gaze. Finally, these direct gaze events were sporadically induced by the experimenters, and it is unknown if they were mixed with other social cues. We do not believe rhesus macaques would leave gaze-following to chance – so what would prompt a rhesus macaque to follow the gaze of another? We suspected they may be able to resort to using direct gaze to initiate gaze-following like humans. Moreover, it seems that macaques are able to distinguish between facial expressions directed towards them versus those that are averted, although this has never been explicitly tested.

Great strides have been made in studying monkey social interactions in natural settings in a quantitative manner. However, monitoring eye movements with sufficient reliability and precision has remained difficult under such circumstances compared to the higher degree of quantification offered by laboratory environments. Studying the interaction of facial expressions and gaze-following – the interest of our study – in unconstrained settings is particularly challenging. The fragile and in most cases unfavorable geometric relationship between the camera and interacting monkeys makes it tremendously difficult to collect high resolution data on interesting combinations of expressions and gaze behavior of the agents. Moreover, much of what we currently know about the effects of direct gaze and facial expressions on gaze-following has been performed in humans, and the few studies done in monkeys are little controlled field studies, exploring these events taking place by chance, or suffer from the limitation that human demonstrators interacted with monkeys. Therefore, in an attempt to explore the interdependence of direct gaze, facial expressions and gaze-following in monkeys in a more rigorous manner and to avoid the pitfalls of unnatural, learnt interspecies interactions, we studied gaze-following responses of monkeys looking at portraits of conspecifics presenting distinct facial expressions while staring at the observer, followed by gaze shifts towards objects. Our results provide clear evidence of the behavioral impact of evaluating danger-related expressions in direct gaze, likely enhancing the observer’s vigilance and preparing their gaze-following response for conflict avoidance.

## 2. Results

### 2.1 Behavioral Paradigm

The behavioral paradigms used in the present study are adapted from the gaze-following paradigm which had previously been used in the discovery, exploration, and causal manipulation of the gaze-following patch (GFP) in macaques (10,18,19). In the first part of this study (Experiment 1), we were interested in the effects of direct gaze and facial expression on gaze-following in macaques (Fig. 1A). The averted head of a conspecific (averted gazer) was presented on a monitor in front of our observing experimental monkeys, and for each trial the observer was required to follow the head gaze of the conspecific towards one of four possible targets presented in a row in front of the conspecific, and make a saccade towards the target singled out by the conspecific’s head gaze. These averted portraits carried a neutral expression. Importantly, prior to the gaze-following portion of the trial, the portrait of the same monkey individual was presented in the center of the screen, in the same location as the averted gazer but here the monkey’s portrait (direct gazer) faced the monkey observer. The direct gazer either looked straight ahead towards the observer (portraying direct gaze) with the eyes open or, alternatively, closed. This portrait could also display a neutral, threatening (open mouth), or fear-grin (silent bared-teeth) expression. All portraits, whether direct or averted, were prepared from photos of the same three monkeys participating in this study. More details of their preparation are included in the Methods section. Additional categories also used as the direct gazer included inverted and scrambled versions of these stimuli (Fig. 1B). These two categories were added as a means of experimental control to disrupt normal face-processing. While the former may elicit the face inversion effect (which will be detailed in section 2.3), the latter preserves low-level visual features but destroys holistic face-processing.

**Figure 1.**
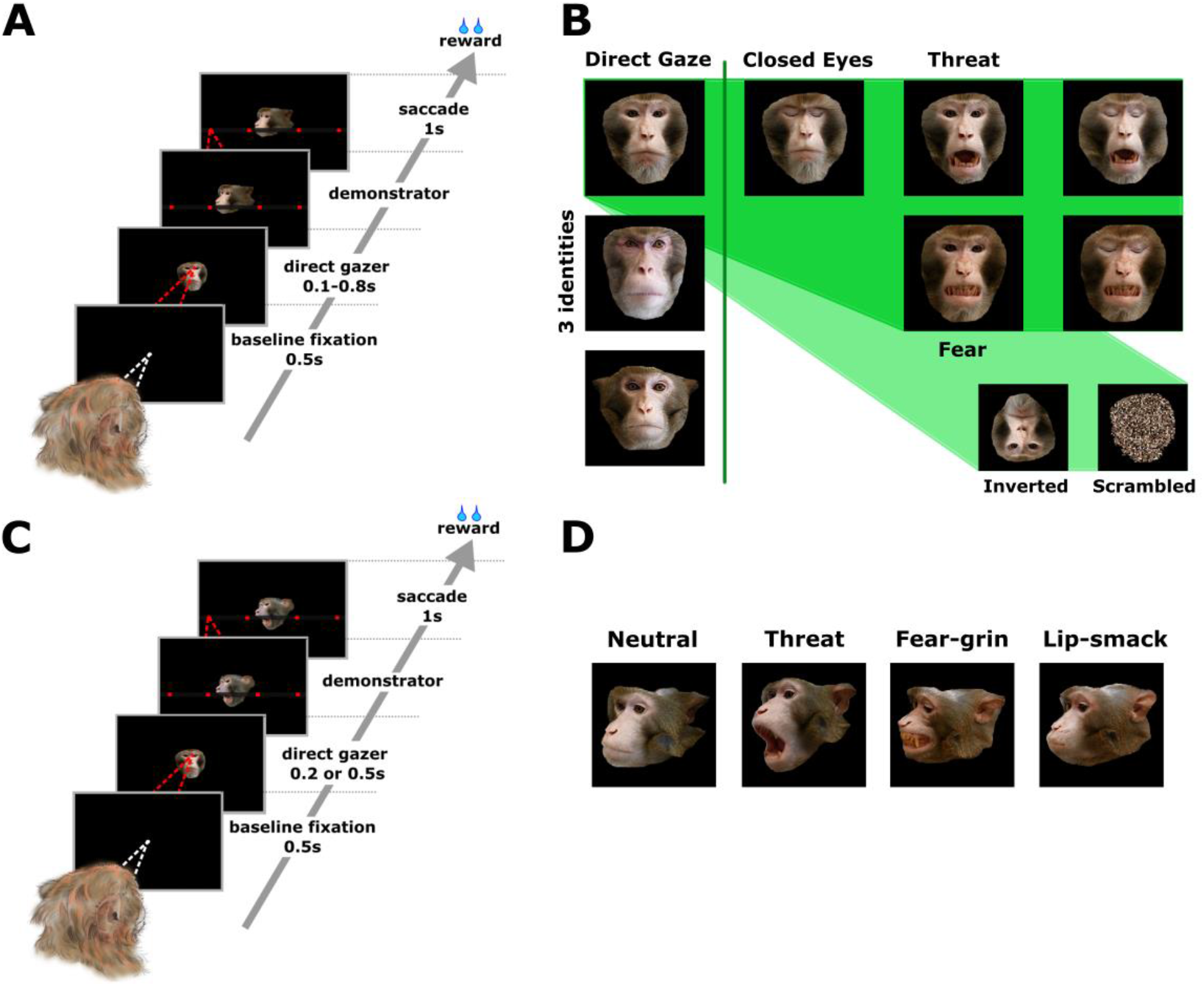
Behavioral Paradigms and Stimuli. **A**. Sequence of events in each trial of the head gaze-following paradigm used in Experiment 1. **B**. Direct gazer stimuli used in Experiment 1. In this visual, we show all the variants of one direct gazer identity, which include his three facial expressions (neutral, threat, and fear-grin) and eyes open and closed conditions, as well as examples of inverted and scrambled stimuli of the same direct gazer in the lower right corner. Variants of the other two identities of the direct gazers are not shown here. **C**. Sequence of events in each trial of the head gaze-following paradigm used in Experiment 2. **D**. Averted demonstrator stimuli harboring facial expressions of one of the monkey identities used in Experiment 2. From left to right, we have a neutral, open-mouth threat, fear-grin, and lip-smacking expression. All four examples shown here are oriented 40° rightward from the demonstrator’s perspective (or 10° leftward for the observer).

In the second part of our study (Experiment 2), we aimed to assess if facial expressions oriented towards the targets had any effect on the latency of gaze-following. The gaze-following portion of the paradigm was modified so that the averted gazing head of the conspecific did not just display neutral expressions, but also threat, fear-grin, or lip-smacking expressions (Fig. 1D). At the time Experiment 2 was prepared we were only able to elicit lip-smacking in one monkey (monkey C) serving as model for the portraits, which is why all lip-smacking demonstrator portraits are based on this particular monkey. For the other expressions (neutral, threat, and fear-grin) all 3 monkeys that took part in this study were able to serve as models. The demands of the task were the same as in Experiment 1, and the observing subject was required to use the head gaze of the demonstrator to single out the correct target (Fig. 1C). Considering the findings obtained in Experiment 1, we restricted the direct gazer type in Experiment 2 to only upright neutral and threat expressions, portraying open or closed eyes, and waived the scrambled versions of the various direct gazer categories.

Trials in both Experiment 1 and 2 commenced with fixation of a central white colored fixation point, and 500ms after trial onset the direct gazer was displayed, centered behind a red fixation point. As described before, the direct gazer consisted of a forward oriented monkey portrait, with his eyes open or closed, and presenting a neutral, threatening, or fear-grin expression. The direct gazer could be presented upright or inverted, and every category of stimuli also had a scrambled equivalent, which was produced via shuffling of the pixels of the portrait in a random manner to disrupt the holistic facial image while preserving low level visual features. The direct gazer in Experiment 1 was presented for 100, 200, 300, 400, or 800ms, while in Experiment 2 only two durations were used, 200 or 500ms, after which the direct gazer disappeared along with the red fixation point, and was replaced by a demonstrator monkey gazing towards one of four spatial targets. For Experiment 1, the direct gazer categories and their different presentation durations were completely randomized (Fig. 1A). In Experiment 2 only one direct gazer category was tested per experimental session (neutral or threatening), as there were four demonstrator gaze directions (i.e. the portrait gazing at one of four targets) each associated with four (neutral, threatening, fear-grin, and lip-smack) facial expressions (Fig. 1C). As control, we used a black background, which was presented for the same durations as that of the portraits mentioned above.

Unlike the gaze-following paradigm used in prior studies, we opted to not use a go cue as it may conceal a potential influence of the direct gazer on the latency of the subsequent gaze-following response. Instead, the appearance of the demonstrator monkey served as a go cue to initiate gaze-following, and the experimental monkey could initiate his gaze-following response whenever he was ready. Identification of the correct spatial target that the demonstrator monkey was looking at was followed by a juice reward. The identity of the demonstrator monkey used as a spatial cue was always consistent with the identity of the direct gazer.

### 2.2 Direct gaze facilitates gaze-following when accompanied with a threat expression

In Experiment 1, we explored whether the expression associated with direct gaze influenced the onset of gaze-following responses. We separated all gaze-following trials according to their direct gazer categories, and identified the average response latency of the gaze-following saccade. The latencies for the control condition (black background) were subtracted from the latencies of the gaze-following saccades of each of the direct gazer categories (and for each viewing duration) to standardize the latency measures.

Figures 2A-I summarize the effects of direct gaze with eyes open (red) and eyes closed (blue) on the latency of the gaze-following response for each type of facial expression (Fig. 2A-C for monkey L, D-F for monkey J, and G-I for monkey C). A 7 × 2 × 5 ANOVA (direct gazer type x eye visibility [direct gaze vs closed] x presentation duration) revealed an interaction between eye visibility and direct gazer type for two of the monkeys (Monkey L: F=2.34, df=6, p=0.0294; Monkey C: F=2.76, df=6, p=0.011), while in a third monkey an interaction between eye visibility, direct gazer type, and presentation duration was found (Monkey J: F=2.94, df=24, p<0.0001). For all three monkeys, direct gaze from the eyes had no significant effect on gaze-following responses when the direct gazers were displaying neutral expressions, for any presentation duration (Fig. 2A, D, and G, posthoc t-tests not significant). This result is contrary to findings in humans, where direct gaze coupled with a neutral expression is sufficient to trigger an earlier gaze-following response (8). However, when direct gaze with eyes open was paired with the threat expression, gaze-following response times were significantly earlier, compared to if the threat expression was associated with closed eyes (Fig. 2B, E, and H). This effect was significant when the threatening direct gazer was displayed for 200ms in monkeys L and J (posthoc t-tests, p=0.0028 and p=0.0065 respectively, using the Benjamini-Hochberg procedure at a false discovery rate of 5%), and for 400ms in monkey C (p=0.0078). We did not find effects for other presentation times for the monkeys, but that is likely because shorter presentation times do not allow sufficient time for the stimulus to be processed, while longer presentation times lead to habituation of the stimulus.

**Figure 2.**
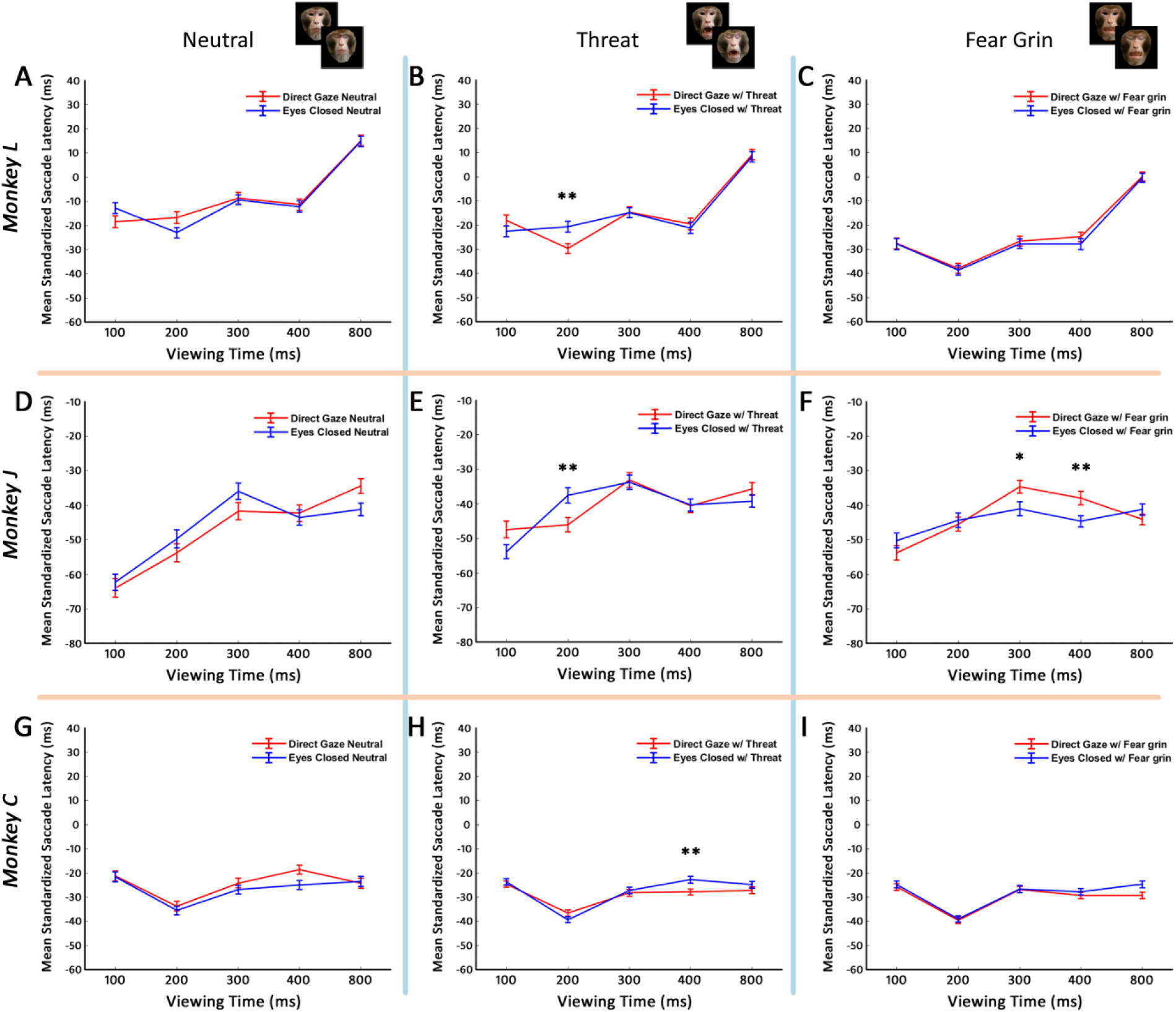
Effect of eye visibility and facial expressions of the direct gazer on gaze-following responses in Experiment 1. Columns depict facial expression types of the direct gazer (from left to right: neutral, threatening, fear-grin). Red and blue lines represent eyes open and eyes closed conditions of the direct gazer respectively. Mean gaze-following responses ± SEM are shown, and posthoc t-tests were performed, ** P<0.01. Benjamini-Hochberg procedure was applied to control for the false discovery rate (0.05). **A-C**. Results for Monkey L: 2B shows significantly faster gaze-following responses for a threatening direct gazer with eyes open compared to the same expression with eyes closed (200ms display time). **D-F**. Results for Monkey J: 2E shows significance of eye visibility when a threatening direct gazer is displayed for 200ms. **G-I**. Results for Monkey C: 2H shows significance of eye visibility when a threatening direct gazer is displayed for 400ms.

In combination with a fear-grin expression gaze-following response times were delayed (Fig. 2F, posthoc t-tests p=0.0174 and p=0.0081 for presentation times 300 and 400ms respectively). However, this effect of delaying gaze-following with a directly gazing fear-grin expression was only observed in monkey J. Our results suggest that not only are monkeys sensitive to the direct gaze of a conspecific and use it to accelerate gaze-following, but they also experience it in a manner different from humans and require reinforcement from facial expressions.

### 2.3 Gaze-following in macaques is affected by face inversion

The face inversion effect, characterized by more rapid recognition and discrimination of upright as compared to inverted faces is a phenomenon observed in humans and chimpanzees (20). There have been many conflicting reports as to whether macaques experience the face inversion effect, because there is no agreed way to test for it in monkeys (21–24). To the best of our knowledge the effect of face inversion on gaze-following is untested in both humans and non-human primates. In Experiment 1, we introduced inverted faces that included various facial expressions, with or without direct gaze, and assessed the effects of these stimuli on the latency of gaze-following. If inverted and upright direct gazers were processed in the same way, we would have expected identical gaze-following profiles. Here we show that is not the case. While the upright directly gazing threat expression had a fairly consistent effect across all our monkeys, each of our monkeys had a unique response profile associated with inverted directly gazing faces. For instance, the gaze-following responses of monkey L were not affected by direct gaze at all when presented in the inverted configuration (Supp. Fig. 1A-C), whilst monkey J displayed earlier gaze-following responses when confronted with inverted expressions with their eyes closed (Supp. Fig.1E and F). Finally, Monkey C even showed earlier gaze-following responses when confronted with inverted neutral faces with their eyes closed (Supp. Fig.1G).

Whatever the reason for these inconsistencies may be, the clear deviation from responses to upright faces suggests that in macaques inverted faces are processed differently from upright faces. Otherwise, the impact of direct gaze from the eyes combined with the threat expression would have been preserved, which we do not observe. On the other hand, we also recognize that inverted faces are artificial configurations; even more uncanny when they rotate back to the upright orientation of the demonstrator, and is unlikely to be a familiar experience for rhesus macaques. This could have also contributed to the profound inter-individual differences in our findings. Finally, direct gaze had no significant effect for scrambled direct gazers, which is what we expected (posthoc t-tests not significant).

### 2.4 Gaze-following performance is not affected by direct gazer category

Given that direct gaze and threat expressions can accelerate gaze-following, we wondered if threat and more generally the particular expression associated with direct gaze also mattered for the accuracy of subsequent gaze-following. Hence, we calculated the gaze-following performance for each of the direct gazer categories independently of the presentation duration and asked if certain direct gazer categories had an impact on the gaze-following efficiency as gauged by the percentage of correct target choices. However, we were unable to uncover any such effects in our monkeys (posthoc t-tests not significant).

### 2.5 Affiliative expressions towards spatial targets delay gaze-following responses

In humans, facial expressions convey different meanings depending on whether they are directed towards the observer or another location of interest (25–27), and we suspected that this may also hold true for macaque monkeys although this has never been explicitly tested. We maintained our original question of whether or not direct gaze could affect gaze-following, but now asked how expressions added to the spatial cue might modulate gaze-following responses (Fig. 1C and D). Given our finding that gaze-following could be influenced by prior direct gaze, we speculated that subsequent facial expressions could have an add-on effect to gaze-following. If we take what is reported in the human literature to also hold for rhesus macaques, one might expect that a fear-grin expression following a threatening direct gaze might entail extremely early gaze-following responses.

The experiments testing this assumption were divided into two groups; sessions that displayed neutral faces as the direct gazer, and sessions that made use of threatening faces as the direct gazer. This reduced the number of possible facial expression combinations with the spatial cue expressions, i.e. the expressions accompanying the oriented faces. To sum up, in a neutral-based session a neutral face direct gazer (with eyes showing direct gaze or eyes closed) was followed by a spatial cue portraying a monkey averting gaze in association with one of four expressions (neutral, threat, fear-grin, or lip-smacking), and likewise for a threat-based session. We also simplified the direct gazer presentation durations to short (200ms) or long (500ms), also for the purpose of lowering the number of potential combinations of direct gazer expression type, gaze type, presentation duration, and spatial cue expression. The reduction of the types of combinations also afforded more repetitions within each session, and therefore could decrease the number of experimental sessions in Experiment 2 (Fig. 1C).

We conducted a 2 × 2 × 4 ANOVA (eye visibility [direct vs closed] x duration (200 vs 500ms] x spatial cue expression [neutral, threat, fear, lip-smack]) for neutral and threat direct gazer sessions. Regardless of which direct gazer type we used, we no longer saw any influence of the direct gazer on the gaze-following response of any of the observing monkeys, nor any consistent interaction between eye visibility and the other factors. This is an unexpected outcome as one might have expected that a threatening direct gazer with his eyes open would continue to facilitate gaze-following. However, there was no difference between gaze-following responses that started with threatening eyes open or threatening closed eyes direct gazers, even when followed by a neutral face gazing towards one of the four targets. It is possible that the addition of the second facial expression in the demonstrator overshadows the information provided by the eyes in the direct gazer, leading to the loss of the latter’s effect on gaze-following.

Under the assumption that the eye visibility of the direct gazer does not matter when followed by an expressive demonstrator, we pooled the data over all eye visibility conditions for neutral and threatening direct gazers and separated the resulting sets according to only direct gazer presentation duration and demonstrator expressions. This also allowed us to double our sample size as we would effectively only be performing a 2 × 4 ANOVA (duration x spatial cue expression) with the same experimental results. We identified an interaction between the averted gaze duration and expression (Monkey L: F=19.63, df=3, p<0.0001; Monkey J: F=4.98, df=3, p=0.0019; Monkey C: F=10.27, df=3, p<0.0001) when the direct gazer was neutral. Importantly, we also detected significant differences between gaze-following responses within the facial expression category (p<0.001 for all 3 monkeys).

Despite the simplification of the paradigm and reduction of the number of configurations we had to compare, it was difficult to conclude how our subjects were reacting to the different presentation durations, direct gazers, and the different expressions on the spatial cues, especially since we were also comparing across three monkeys. Our statistical tests informed us that the facial expression category of the spatial cue caused significant differences in gaze-following latencies, but at first glance it was difficult to draw a meaningful conclusion as to how the expressions were affecting gaze-following. We decided to normalize and pool all gaze-following responses as a first step towards interpreting our data. Fig. 3A and B show results collected from sessions that utilized neutral direct gazers and threat direct gazers respectively. We found that in sessions that began with a neutral direct gazer (Fig. 3A), gaze-following responses resulting from spatial cue expressions that were affiliative (fear-grin and lip-smack) were significantly later than those that were neutral or antagonistic (threat). This result was the same for both direct gazer presentation durations.

**Figure 3.**
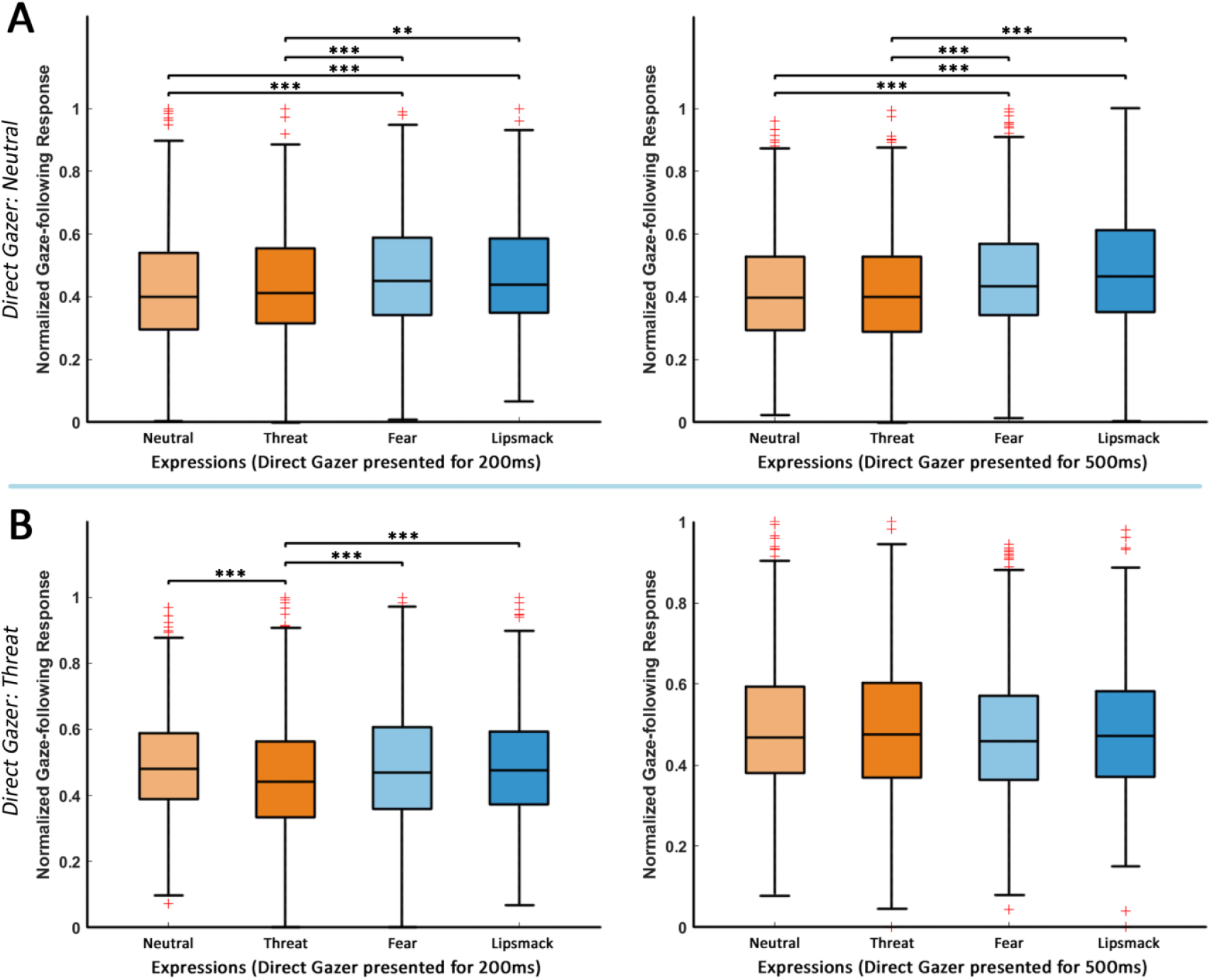
Effect of demonstrator expression type on gaze-following responses in Experiment 2. Because eye visibility no longer mattered in Experiment 2, we pooled together all trials portraying neutral (A) and threatening expressions (B) regardless of eye visibility respectively. Gaze-following responses per subject for each demonstrator expression category were then normalized between 0 and 1, and then pooled together. The solid black line within the box represents the median normalized response, and the whisker lengths are set as 1.5x the interquartile range. Red crosses represent 1.5x outliers. **A**. Left and right plots show normalized responses where the direct gazer expression was neutral and presented for 200 and 500ms respectively. Gaze-following of affiliative expressions is significantly slower than for antagonistic expressions, regardless of direct gazer presentation duration. **B**. Left and right plots show responses where the direct gazer expression was threatening and presented for 200 and 500ms respectively. Gaze-following of the threat expression is faster than the following of the other facial expressions, but only when the prior threat direct gazer was presented for 200ms.

In the sessions that were initiated with a threatening direct gazer (Fig. 3B), the spatial cue with the threat expression facilitated gaze-following responses the most (left). Unlike the result with neutral direct gazers where the gaze-following responses were divided between affiliative or antagonistic expressions, now the response for the threatening demonstrator stands out the most among the four expressions. It is possible that its consistency with the expression of the direct gazer prevents the dilution of the social salience of the cue, whereas in the other conditions an expression change occurs, i.e. from threat to lip-smacking, which leads to the influx of new expression information. What is apparent is that in both direct gazer conditions affiliative expressions consistently lead to later gaze-following responses. This effect is not restricted to specific presentation durations, and this is likely due to the appearance of the second expression in the spatial cue overriding all effects of the direct gazer. Taken together, the results of Figures 2 and 3 suggest that while direct eye gaze can be a powerful cue in expediting gaze-following when combined with a threatening expression, the presentation of new information will take priority and determine the gaze-following response, irrespective of the direct gazer features. Where an antagonistic face is looking at seems to take precedent over where an affiliative face is attending to.

### 2.6 Direct expressions lead to earlier gaze-following expressions than averted expressions

In this final part, we wanted to compare gaze-following responses from Experiments 1 and 2. Previously, we had separately explored the effects of facial expressions when they were directed towards the observer and when they were averted and directed towards spatial targets. This presented a unique opportunity to assess within the same experimental monkeys if expressions were perceived differently depending on where they were directed, using their gaze-following responses as a readout. Past research had shown that monkeys may be more likely to follow gaze if an averted face was expressing fear (13), but it is unknown how that same monkey would have reacted if the fear expression had been directed towards himself prior to gaze-following. By combining the results of Experiment 1 and 2, we explored the impact of different expression transitions from direct to target oriented gaze (direct gazer to demonstrator) on the observer’s gaze-following. We only used trials that had direct gazer presentation durations of 200ms because that was the only timing shared between both experiments.

The results for the three monkey individuals are summarized in Fig. 4A-C. As a reference, we used trials from both Experiments 1 and 2 in which the neutral expression was maintained from the direct gazer to the spatial cue (column 1 in each plot). The figures show a clear and significant delay in gaze-following whenever there was an averted expression. In two of our three monkeys this effect was present for both threat and fear expressions (two sampled t-test, monkey L: p=0.00221 and p<0.001 respectively; monkey C: p<0.001 for both), whereas the third monkey (Fig.4B) only showed a significant effect for the fear expression (p<0.001). One interpretation is that the observing monkeys place less importance on expressions that are not directed towards them, and therefore follow gaze after a longer pause. Alternatively, the demonstrator expressions (which appear later in the course of each trial) may exert an increased informational load on the observer, causing delayed gaze-following responses.

**Figure 4.**
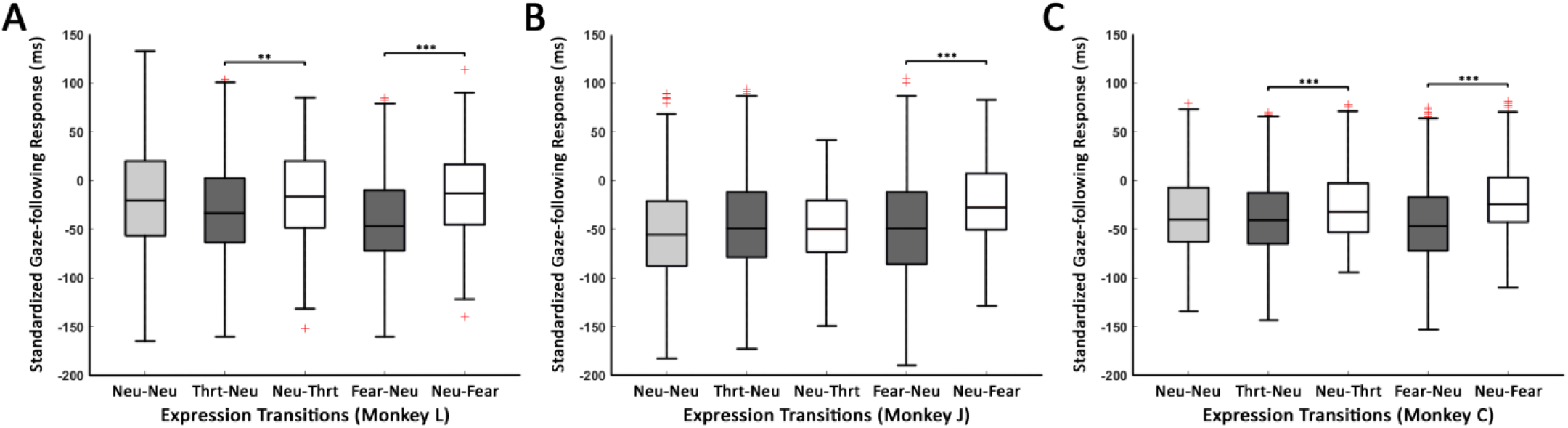
Comparing the effects of facial expressions on gaze-following depending on whether the expression was directed towards the observer or the spatial target. The solid black line within the box represents the median, and the whisker lengths are set as 1.5x the interquartile range. Red crosses represent 1.5x outliers. **A-C**. Results from monkeys L, J, and C respectively. Two sample t-tests were performed, ** P < 0.01, *** P < 0.001. Abbreviations: Neutral (Neu), Threat (Thrt). Neu-Thrt signifies a transition from a neutral direct gazer to a threatening demonstrator, and vice versa for the other notations. Expressions that were portrayed in the demonstrator significantly reduced gaze-following reaction times compared to the same expression that was directed towards the observer as a direct gazer; for A and C both threat and fear, and only fear in B.

## 3. Discussion

When the other’s eyes meet our own, we experience direct gaze, a potent visual cue indicating that we have become the focus of the other’s attention. This is a vital step towards establishing mutual gaze, allowing both parties to silently acknowledge their attention to each other. If the other subsequently looks towards a third object, the observer may be compelled to follow the other’s gaze to the same location, thereby establishing joint (object-related) attention. Gaze-following may also occur without prior gaze, yet at least in humans it may be promoted by preceding direct gaze (8). Whether monkey gaze-following also benefits from direct gaze in the same manner was unclear. Hence the primary aim of the present study was to test whether monkeys are able to use direct gaze to inform their gaze-following decisions and, moreover, to determine the impact of accompanying facial expressions.

The first experiment’s results clearly indicate that direct gaze can indeed enhance gaze-following responses in monkeys, yet only when accompanied by an expression of threat. However, the synergistic effect of direct gaze and threat is not independent of the demonstrator’s expression when shifting gaze to a target. Our second experiment showed that whenever a distinct facial expression accompanied the demonstrator’s gaze aversion, neither the eyes nor the expression of the direct gazer mattered anymore. When the expression accompanying gaze aversion indicated submission, the gaze-following response was delayed compared to both neutral and threatening expressions. Finally, direct comparison of both experiments revealed that facial expressions accompanying direct gaze triggered earlier gaze-following responses than the same expressions accompanying the demonstrator’s gaze shifted towards a target (summarized in Table 1).

**Table 1.**
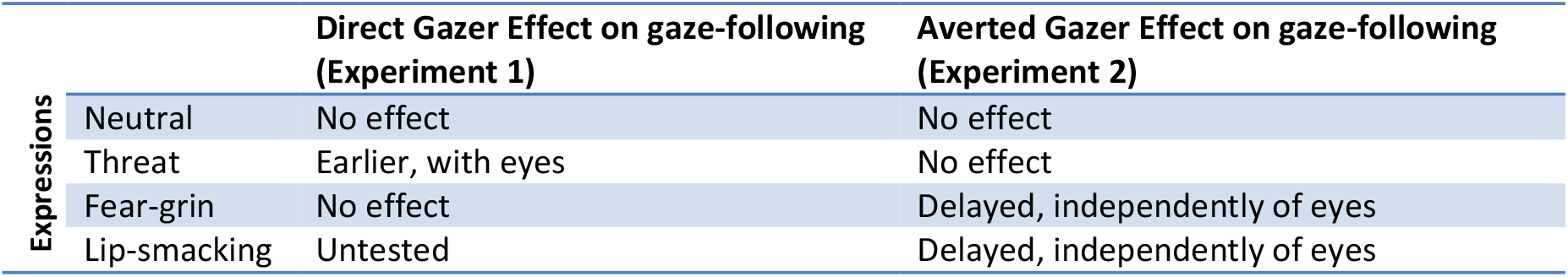

In order to avoid the many shortcomings of studying social interactions in natural settings outlined in the introduction, we resorted to a study of the interaction between gaze-following and facial expressions under laboratory conditions guaranteeing stable high resolution recordings of the observer’s eye movements prompted by exposure to demonstrator portraits offering combinations of highly standardized facial expressions and gaze orientations from an extensive library. In an attempt to standardize the demonstrator stimuli as much as possible we used static stimuli rather than video clips. The ease of interpreting our results in terms of ethological validity promotes trust that our paradigm indeed reflects natural social behaviour of macaques. If these views are truly justified we must await ground truth studies with comparable data quality that may one day become feasible also under natural conditions.

In the wild, facial displays of threat and aggression accompanying direct gaze are frequently employed by dominant males to assert their status within their group and to maintain control over resources (11,28). It would be extremely beneficial for others in the group to pay attention to these signals in order to get ready for the potential consequences of a dominant male’s upcoming move as this could be hostile and potentially dangerous. As these moves are typically initiated by gaze shifts to third parties or objects of mutual interest, the observer – no matter if competitor or subordinate – would be well advised to follow gaze as quickly as possible. Findings by Champ et al. (17) showed that appeasing and submissive behavior attracted more gaze-following from an observer, which is at odds with what our results show. The reason for this discrepancy might be that in our study each gaze shift of the demonstrator was preceded by an exchange of direct gaze, a strict sequence that was lacking in the Champ study.

The existence of a face inversion effect in monkeys has been subject to much debate: some studies have reported no change in face recognition upon inversion of faces (21,22), whilst others have reported an inversion effect in the sense of an impaired ability to recognize inverted faces (24). Another study even reported an inversion effect for human but not for monkey faces (23). If inverted faces are not processed in the same way as upright faces (i.e. inversion effect present), one might expect threat expressions combined with direct gaze to have a similar effect on gaze-following no matter of the orientation of the face. Our data shows that this is not the case, as inverting the direct gazer exhibiting a threat expression saw the loss of effect of direct gaze on the subsequent gaze-following response that differed between individuals. The lack of consistency may reflect individually differing perceptual interpretations of inverted faces. After all, instances in which a monkey might encounter a fully inverted conspecific exhibiting direct gaze in association with meaningful expressions must be rare, if not completely absent. In other words, such configurations are simply unnatural and if presented in an experiment may prompt highly idiosyncratic interpretations.

In Experiment 2, the facilitatory influence of threat accompanying direct gaze on gaze-following was cancelled as soon as a new averted facial expression appeared. The reason may be that the appearance of two expressions in quick succession created a redundancy of social signals, leading to the prioritization of the more recent expression and therefore the loss of influence of direct eye gaze. It is in the best interest of monkeys to respond quickly to social cues and changes in their environment for survival, which is why the later expression with the gaze shift may take precedence. Recency bias is a documented phenomenon in macaques (29), although it has not been explored in the context of social cue retention. Finally, also restrictions of processing capacities seem conceivable, entailing an attentional focus on the second facial expression, especially when this is necessary in our paradigm for reward.

In monkey groups, gaze-following aids in the determination of rank with the dominating individual receiving the most attention, often associated with signs of submission. Following the gaze of group members expressing submission helps others to shift attention to this dominating and potentially dangerous individual (30). As described above, the results of Champ et al. (17) support the view that this shift is facilitated by concomitant fear-grin as an indicator of submission to a higher ranking individual. Similarly also the work by Goossens et al. (13) showed more gaze-following responses when a gaze-shift was accompanied by fear-grin. Against the backdrop of this work, one might have expected that also in our study the demonstrators exhibiting fear-grin in conjunction with their shifting gaze would elicit earlier gaze-following responses. Instead, the responses to the demonstrator’s averted gaze, associated with fearful or lip-smacking expressions actually occurred later (Fig. 3A and B) in our study. Hence, how can we reconcile this difference?

In humans, averted expressions of fear have been found to be difficult to disengage from, leading to weaker gaze-cueing effects, especially if the subjects were prone to anxiety (31–33). Surprisingly, averted rather than direct gaze has been found to elicit increased arousal in monkeys (34). A directly gazing face indicates the attention of the other is on the observer, and while this is socially significant for macaques, they may have adapted to downregulate their level of arousal, for instance by breaking eye contact. However, an avertedly gazing face points to a yet unknown target of interest which needs to be identified, adding an element of unpredictability which demands vigilance and may contribute to the increase in physiological arousal. Combined with the inherent ambiguity of the fear-grin expression which may indicate anxiety/fear rather than submission, we conclude that the need to closely analyze arousing averted expressions conveying important information on potential danger may be the cause of the delayed attentional disengagement and therefore later gaze-following responses.

The fact that delayed gaze-following responses occur only when facial expressions are accompanying the conspecific’s gaze shift demonstrates that rhesus macaques can flexibly interpret expressions based on context analogous to humans. Modulation of gaze-following and the gaze-cueing effect via changes in context provided by facial expressions has been recognized in humans, although the diversity of paradigms used makes it difficult to pinpoint what exactly the interaction is between gaze direction, eye visibility, and facial expressions (35). By contrast, the way the perception of an expression can change based on gaze is a more consistently reported phenomenon in humans. According to the shared signal hypothesis gaze direction influences the perception of facial expressions when they communicate similar intent (36). For instance expressions such as joy or anger are perceived as approach-type expressions and are therefore enhanced when viewed directly, whereas avoidance-type expressions like sadness and fear are promoted by averted gaze (25,26). If we break down both of our experiments in terms of events that might occur in the wild, it seems there is more urgency for an observing monkey to respond to aggressive intent from a conspecific than to identify what the conspecific may be threatening in the environment. In the same vein, monkeys seem to prioritize interpreting another’s fear in them rather than discovering what may have frightened the other.

While humans can demonstrate enhanced gaze-following in response to direct gaze alone (8), we were unable to observe a comparable result in any of our macaques. A factor contributing to this discrepancy may be the differential processing of social signals that hinges on our available social cognitive capabilities and how our social groups are structured. The cognitive processes involved in perceiving and responding to facial expressions, particularly in the context of gaze-following, could be shaped by species-specific adaptations. Unlike humans who have evolved a variety of ways to communicate our intentions, monkeys have a much smaller social arsenal to draw upon, and may have adapted responses that prioritize the detection and response to potential threats, and to avoid meaningless conflict (37– 39).

Sensitivity towards direct gaze is a phylogenetically ancient feat not confined to primates, and has been discovered in other mammals, and even some species of bird and fish (40). Because the direct gaze of predators represents danger to the observer, it was beneficial for many animals to develop the ability to detect direct gaze. Rhesus macaques have been rated high in their tendency to use direct gaze as a threat signal and are reluctant to make prolonged eye contact (11). In other words direct gaze is generally not regarded as an affiliative gesture. By contrast, gaze-following is recognized as a more evolved cooperative behavior to afford living in a social group (2), arguably derived from the ability to exploit direct gaze. The ability to produce and read facial expressions appeared much later, and for the most part seems to be restricted to the mammalian lineage (41,42). Finally in species prone to aggression, the ability to bridge from a provocative stare into cooperative communication may have been an important step towards establishing social harmony. Our demonstration that rhesus macaques can use direct gaze and the integration of facial expressions to shape gaze-following behavior and modulate its exigency is congruent with this proposal. It would be interesting to examine whether or not possession of a more sophisticated facial muscle control architecture that allows for a wider repertoire of facial movements drives the emergence of more prosocial and affiliative behaviors in other non-human primates and mammals.

## Author Contributions

I.C. and P.T. designed research; I.C. performed research and analyzed the data; I.C. and P.T. wrote the paper and contributed to discussing analysis.

## 4. Materials and Methods

### 4.1 Animals and Surgery

Three male rhesus macaques (*Macaca mulatta*) were involved in the present study. They had all been used before in unrelated electrophysiologcal work. The three – Monkey L (18 years old, 11kg), Monkey J (21 years old, 16kg), and Monkey C (16 years old, 11kg) – were all housed separately and alone. They all had originally played the dominant roles in their previous pairings (all male). Monkey L and J had lost their partners due to natural causes, while monkey C lost his dominant status in his pairing and subsequently had to be separated from his partner to avoid injury. They had never undergone trial pairings nor had they been involved in skirmishes with each other, but had occasional visual contact with each other in the experimental setups and animal facility. While we cannot rule out their familiarity with each other’s faces, we do not believe the monkeys had an established hierarchy as they did not have prolonged contact with one another.

All three monkeys had been implanted with a titanium head-post for restraining the monkey’s head in order to allow precise measurement of eye movements and chambers for electrophysiological recordings from the superior temporal sulcus, the latter not related to the present study. Being able to head fix the monkeys was essential for the quantitative characterization of gaze-following. Surgeries were carried out under combination anesthesia with isoflurane (1.3%) and remifentanil (1 – 2 μg per kg per minute), with meticulous monitoring and control of body temperature, heart rate, blood oxygen saturation, and blood pressure. Opioid analgesics (buprenorphine) were administered until the monkeys showed no signs of residual pain and were given ample time for full recovery before entering the behavioral training for the experiments that preceded the behavioral study at issue here. All experimental preparations and procedures were sanctioned by the local animal care committee (Regierungspräsidium Tübingen, Abteilung Tierschutz), fully complying with German and European law and the National Institutes of Health’s Guide for the Care and Use of Laboratory Animals.

Table 2 summarizes some potentially expedient behavioral observations made on the three monkeys that participated in this study. These observations were collected by the experimenters and the primary animal caretaker at the start of the study.

**Table 2.**
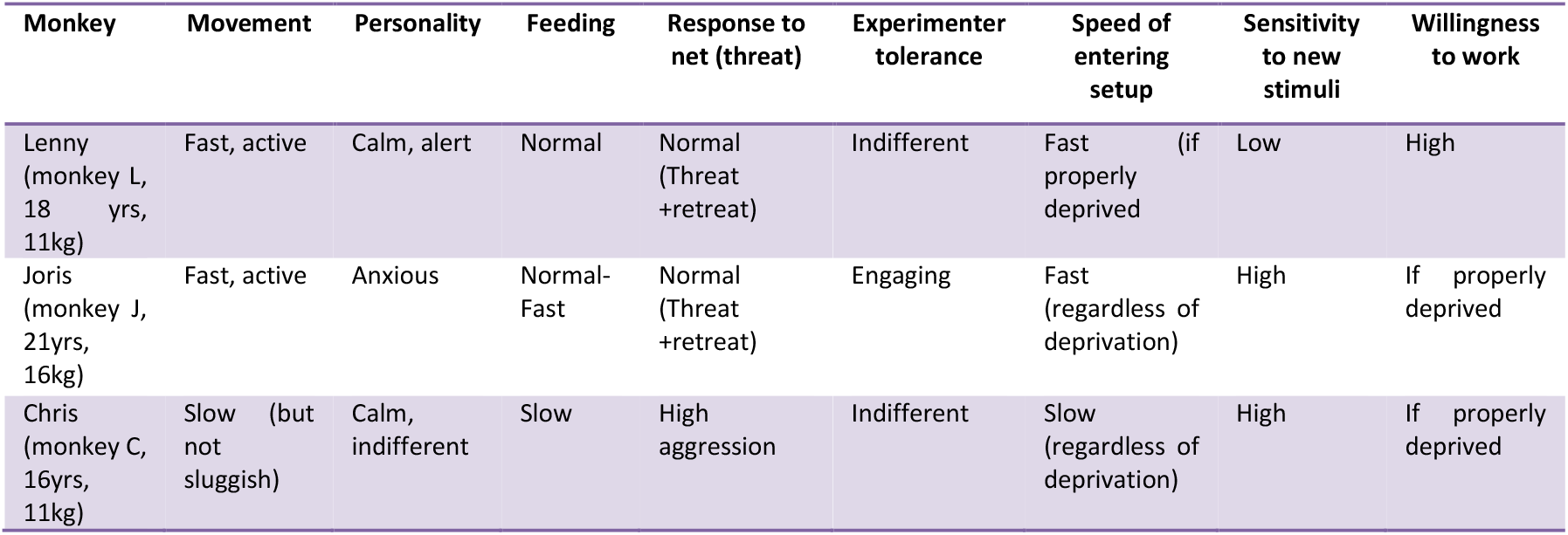

| Monkey | Movement | Personality | Feeding | Response to net (threat) | Experimenter tolerance | Speed of entering setup | Sensitivity to new stimuli | Willingness to work |
| --- | --- | --- | --- | --- | --- | --- | --- | --- |
| Lenny (monkey L, 18 yrs, 11kg) | Fast, active | Calm, alert | Normal | Normal (Threat +retreat) | Indifferent | Fast (if properly deprived) | Low | High |
| Joris (monkey J, 21yrs, 16kg) | Fast, active | Anxious | Normal-Fast | Normal (Threat +retreat) | Engaging | Fast (regardless of deprivation) | High | If properly deprived |
| Chris (monkey C, 16yrs, 11kg) | Slow (but not sluggish) | Calm, indifferent | Slow | High aggression | Indifferent | Slow (regardless of deprivation) | High | If properly deprived |

### 4.2 Behavioral Paradigms

For the present study, the three monkeys were trained on two variants of a gaze-following paradigm that both required them to use head gaze information provided by the portrait of a conspecific demonstrator. Of the three monkeys used in this study, only Monkey L had previously participated in a gaze-following study and therefore was already familiar with the demands of the paradigm. Monkey J and L were new to the gaze-following paradigm. In the final behavioral paradigm, each individual trial began with a white fixation point on a black background, and the experimental monkey was required to fixate on the point for 500ms. Subsequently, the white fixation point was replaced by a red fixation point centered on a portrait gazing at the observer, a configuration we refer to as the “direct gazer” stimulus. In Experiment 1 the direct gazer was presented for 100, 200, 300, 400, or 800ms, while in Experiment 2, the direct gazer was seen for 200 or 500ms. The direct gazer was followed by the presentation of the “demonstrator”, a photograph of the same individual as before, but now with his head gaze turned to a distinct spatial target, randomly chosen from a set of 4 dots arranged on the horizontal axis. Although both the demonstrator and all potential targets remained present until the end of the trial, the observer was encouraged to make a saccade towards the target singled out by the demonstrator’s head gaze as a speedy gaze-following response allowing earlier access to reward. If the observer had successfully initiated the trial by fixating the white central fixation point and correctly followed the demonstrator’s gaze to the correct spatial target, he received a drop of juice as reward. If central fixation failed or the observer’s gaze did not meet the target within 300ms after the appearance of the demonstrator, the trial was terminated and the monkey received no reward. The presentation duration and direct gazer category was randomized from trial to trial. In Experiment 1, 32 sessions were collected in monkey L, 40 in monkey J, and 66 in monkey C. In Experiment 2, 10 sessions were collected in monkey L, 10 in monkey J, and 11 in monkey C. Each session for both Experiments 1 and 2 contained an average of 800 gaze-following saccades. Data for Experiment 1 was collected in approximately 3 months with a pause in between for analysis, before starting Experiment 2 which required around 2 months.

The size of the direct gazer and demonstrator portraits were 5.6° by 5.6°, while the white fixation point and the red spatial targets had a diameter of 0.8° each. The spatial targets were arranged in a virtual horizontal row 1° below the center of the demonstrator portrait. With respect to the observing monkey subject, the horizontal eccentricities of the targets were -10°, -5°, 5°, and 10° (or -40°, -20°, 20°, and 40° with respect to the demonstrator). Tracking of eye movements was accomplished using an infrared camera (iViewX, SensoMotoric Instruments, spatial resolution of <0.3° at 50 Hz).

### 4.3 Stimuli preparation

The demonstrator portraits we used in our paradigms for both the direct gazer and the demonstrator were stills taken of the same three monkeys (L, J, and C) that participated in the study. Video-photography was captured while our monkey subjects were seated in their chairs, but were not head fixed to maximize comfort, ease of moving their mouths as required in 3 of the 4 expressions, and for turning their heads. In Experiment 1, the direct gazers could exhibit a neutral, threat, or fear-grin expression, while in Experiment 2 lip-smacking was added. The neutral expression consisted of a closed and relaxed mouth. It is the default expression of monkeys, present during rest or indifference, not prompting specific behavioral reactions. The threat expression is also known as the open-mouthed threat, in which the mouth is opened and the canines are exposed. Such a display may be used to intimidate other members of a social group, and when displayed between two similarly ranked individuals may be interpreted as a physical challenge. The fear expression is often referred to as the fear-grin, exposing all the teeth while the jaws remain closed and the lips are retracted. Such an expression is not only used to indicate submission, but can also be used as a friendly gesture to signal affiliation and appeasement. Finally, the lip-smacking expression consists of a closed mouth and puckered lips, and like the fear-grin it is categorized as an appeasing expression used to alleviate tension. However the lip-smack is more affiliative, and can be used to signal friendliness rather than subordination.

The expressions documented by the portraits were prompted by the caregivers performing in front of the monkeys in appropriate manners. All video-photography were captured while our monkey subjects were seated in their chairs, but were not head fixed to maximize comfort, ease of moving their mouths as required in 3 of the 4 expressions, and for turning their heads to document averted gaze in conjugation with distinct expression. Neutral expressions were the easiest to capture as they were the default expressions of our monkeys easily produced in calm, non-stimulating situations. Threat expressions were more likely to be produced if a human experimenter approached them too closely, or made abrupt movements. Bringing the video camera close to the monkey was also an easy way to cause them to react with the threat expression, possibly as a proximity warning towards the experimenter. The fear-grin was a rare occurrence in our setup, but the use of a handheld mirror to show the monkeys their reflection, or special face masks with prints of monkey mouths could sometimes induce the fear-grin expression. Finally, the lip-smacking expression could be induced by exposure of our chaired monkey to unfamiliar and unrestrained monkeys in the animal facility. In this setting, the fear-grin expression could sometimes be observed. Because the monkeys habituated quickly to the presence of other monkeys or the various props that we showed them, it was important that all videos were taken before the monkeys got bored and regressed to exhibiting neutral expressions. All monkeys produced the neutral, threatening, and fear-grin expressions, while only one monkey (monkey C) produced the lip-smacking expression. After the videos were taken, we isolated frames in which the monkey was conveying the desired expression with direct or averted gaze and cropped out the background, their bodies, and their implants.

The paradigms used in Experiments 1 and 2 only differed in the types of stimuli used and the presentation duration of the direct gazer stimuli; the structure of each trial was essentially the same. Table 3 summarizes the different stimuli involved in Experiments 1 and 2, as well as the presentation durations used. In some trials, the orientation of the direct gazer portraits could also be inverted. Moreover, scrambled direct gazers were used for additional control in Experiment 1, and as expected there was no effect of eye visibility in scrambled stimuli. Generation of scrambled stimuli was achieved by taking the upright form of the direct gazer, dividing it into evenly sized squares with sides 9 pixels in length and rearranging the squares randomly. This disrupted the facial information whilst retaining the low level visual features of the portraits.

**Table 3.**
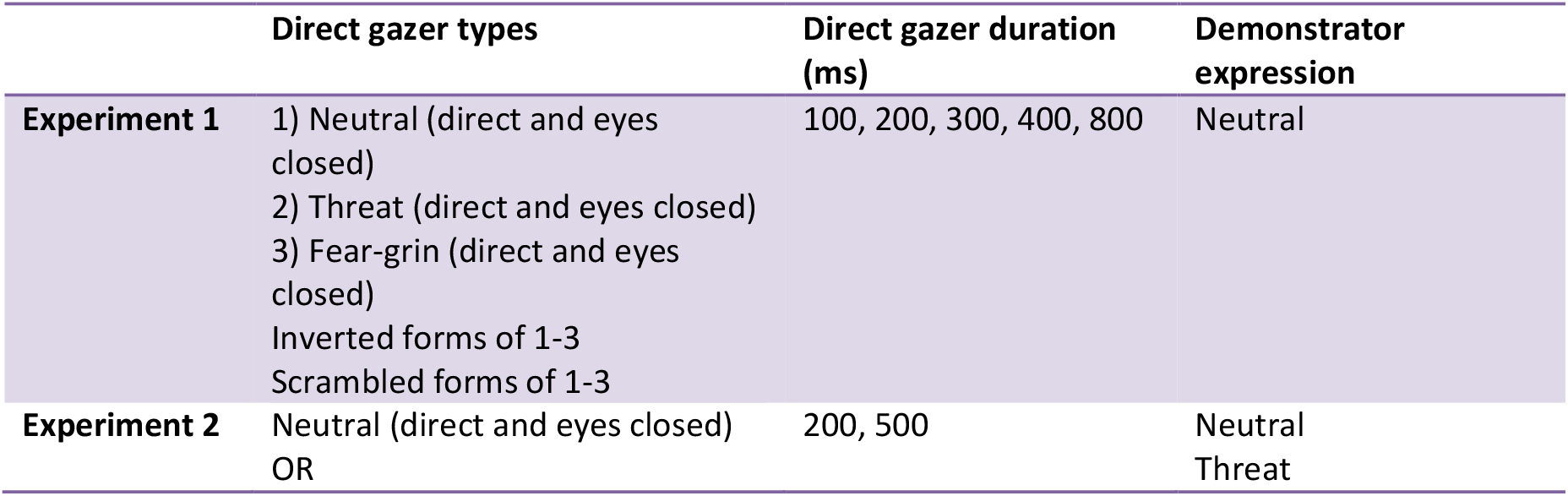
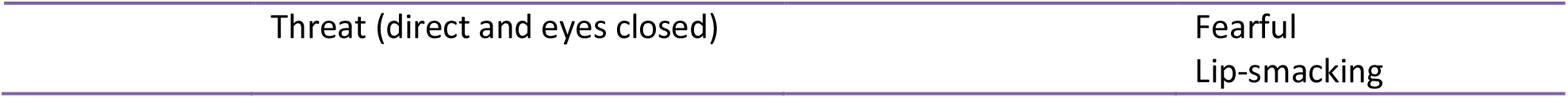

### 4.4 Statistical Analysis

Each daily session of behavioral testing in Experiment 1 consisted of hundreds of gaze-following saccades, but given the number of conditions that were being tested (direct gazer, eye visibility, and presentation duration), the number of saccades made for each condition was few. Therefore, we pooled all the saccades made for each condition across all sessions for each individual monkey for analysis (for example in Monkey L, approx. 12 saccades per category for 75 total stimulus categories totals 900 trials per session, 32 sessions results in 384 trials per stimulus category (gaze-following responses represented by saccadic latencies)). For each trial, we extracted the onset of the gaze-following saccade with respect to the appearance of the spatial cue, and determined the average saccade onset across all sessions for that specific condition (eg. direct gazer: upright threat; eyes: direct; duration: 200ms). Gaze-following saccade onsets were then standardized relative to the average saccade onset produced towards a black background in place of a direct gazer before the appearance of the demonstrator.

To investigate if one of the main variables we manipulated in Experiment 1 or any of their interactions had an impact on the gaze-following saccade latency we resorted to a 7 × 2 × 5 ANOVA (direct gazer type x eye visibility [direct vs closed] x presentation duration), separately for each monkey. Post hoc t-tests were then performed between direct gaze and closed eyes conditions, for all presentation durations, and within each facial expression type. This approach allowed us to uncover which combination of facial expression, presentation duration, and gaze type could modulate the gaze-following response. The Benjamini-Hochberg procedure was applied to control the false discovery rate (0.05).

Since we did not find any effect of direct gaze on the subsequent gaze-following saccade latencies in Experiment 2, regardless of the expression accompanying direct gaze being neutral or threatening or the eyes open or closed, we decided to ignore the eye visibility distinction in order to increase the statistical power of the analysis of the impact of the facial expression of the averted gazer on gaze-following. In other words, in Experiment 2 trials were grouped according to the facial expression of the direct gazer (neutral or threat), presentation duration (200 or 500ms), and the facial expression of the averted gazer (neutral, threat, fear-grin, or lip-smacking). Furthermore, in order to be able to pool data from all three monkeys, we normalized their gaze-following between a range of 0 and 1. A 2 × 4 ANOVA (presentation duration x expression) was then performed on this pooled data for the normalized gaze-following response times, separately for sessions involving a neutral or threatening direct gazer. Posthoc t-tests were performed on groups that were significantly different from each other, and again the Benjamini-Hochberg procedure was applied to control the false discovery rate (0.05).

**Supplementary Figure 1.**
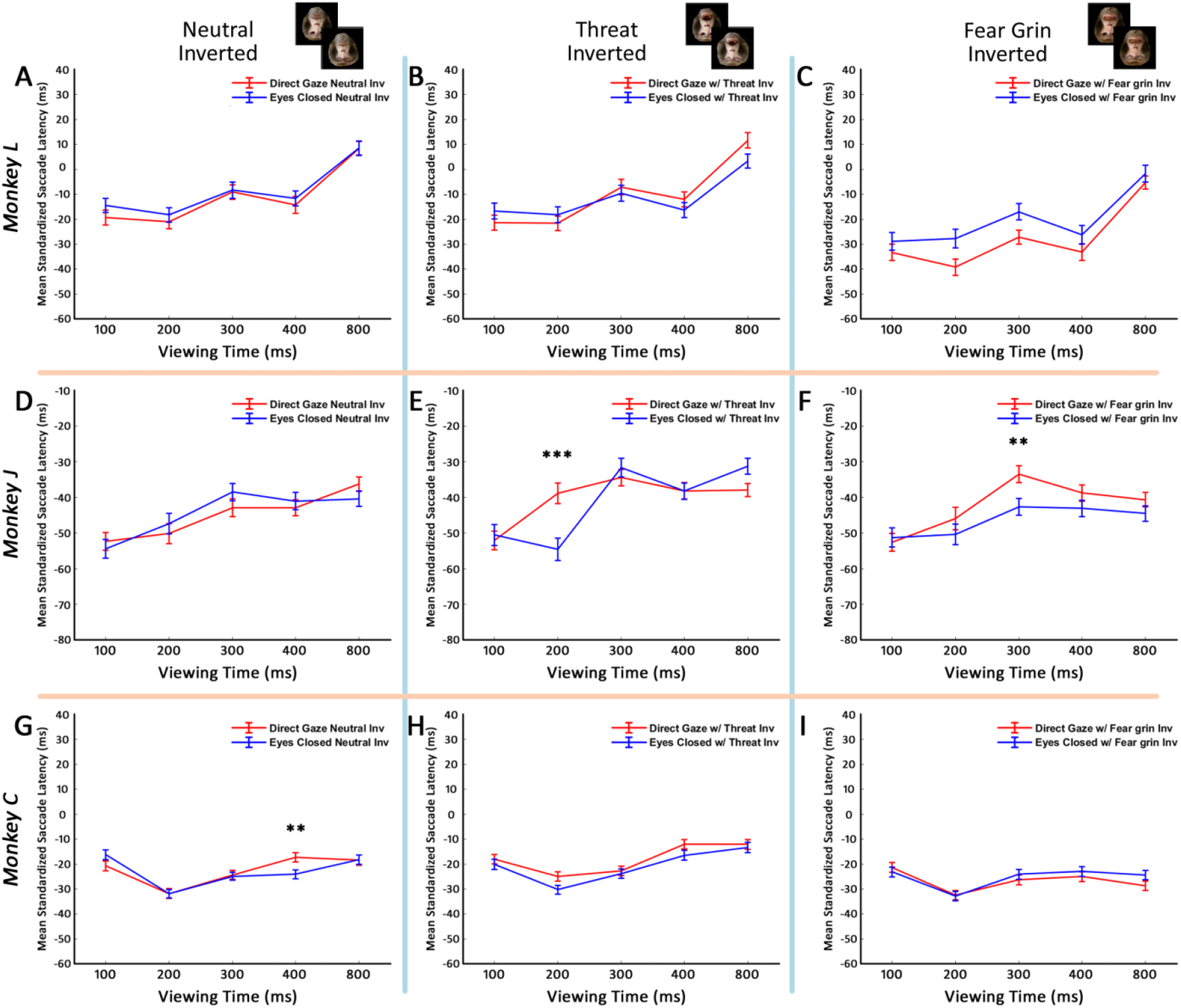
Effects of face inversion of the direct gazer on gaze-following responses. Columns depict facial expression types of the direct gazer (from left to right: neutral, threatening, fear-grin, in their inverted orientations). Red and blue lines represent eyes open and eyes closed conditions of the inverted direct gazer respectively. Mean gaze-following responses ± SEM are shown, and posthoc t-tests were performed, ** P<0.01, *** P < 0.001. Benjamini-Hochberg procedure was applied to control for the false discovery rate (0.05). **A-C**. Results for Monkey L, no significance of eye visibility for any expression. **D-F**. Results for Monkey J: faster gaze-following responses when confronted with inverted threat and fear expressions with their eyes closed (E and F). **G-I**. Results for Monkey C: faster gaze-following responses when confronted with inverted neutral faces with their eyes closed (G).

